# Determination of human DNA replication origin position and efficiency reveals principles of initiation zone organisation

**DOI:** 10.1101/2021.11.30.470574

**Authors:** Guillaume Guilbaud, Pierre Murat, Helen S. Wilkes, Leticia Koch Lerner, Julian E. Sale, Torsten Krude

## Abstract

Replication of the human genome initiates within broad zones of ~ 150 kb. The extent to which firing of individual DNA replication origins within initiation zones is spatially stochastic or localised at defined sites remains a matter of debate. A thorough characterisation of the dynamic activation of origins within initiation zones is hampered by the lack of a high-resolution map of both their position and efficiency. To address this shortcoming, we describe a modification of initiation site sequencing (ini-seq) based on density substitution. Newly-replicated DNA is rendered ‘heavy-light’ (HL) by incorporation of BrdUTP, unreplicated DNA remaining ‘light-light’ (LL). Replicated HL-DNA is separated from unreplicated LL-DNA by equilibrium density gradient centrifugation, then both fractions are subjected to massive parallel sequencing. This allows precise mapping of 23,905 replication origins simultaneously with an assignment of a replication initiation efficiency score to each. We show that origin firing within initiation zones is not randomly distributed. Rather, origins are arranged hierarchically with a set of very highly efficient origins marking zone boundaries. We propose that these origins explain much of the early firing activity arising within initiation zones, helping to unify the concept of replication initiation zones with the identification of discrete replication origin sites.

## Introduction

The replication of eukaryotic genomes requires multiple DNA replication origins that become active at different times during S-phase. However, their position and firing probability remains a matter of debate (1–3). In the late 1960s, visualisation of replication tracts in mammalian cells revealed that several replication origins could be simultaneously active along chromosomes (4), but their position on the genome could not be mapped. It took 14 years to characterise the first eukaryotic initiation site by restriction digestion and autoradiography (5), and a further 24 years to achieve the first fine mapping of hundreds of human replication origins by microarray hybridisation of enriched RNA-capped short nascent strands (SNS) (6).

Early analyses of viral, bacterial and mitochondrial genomes revealed the existence of nucleotide substitution asymmetry that has been linked to replication fork polarity and used to infer the position of replication origins (7–9). By computing the excess of G over C and T over A along one DNA strand, sharp changes from negative to positive values can be seen at the site of replication origins (the so-called Skew-jump or S-jump). Analyses of the human genome also revealed an asymmetry that is attributed to DNA replication and identified 1,012 S-jumps (10, 11). In between two S-jumps, a linear decrease in skew is observed, forming an N-shaped pattern (12). These ‘N-domains’ cover up to a quarter of the human genome. Nevertheless, it is not possible to conclude whether S-jumps result from the firing of a single and efficient origin, or of several less efficient origins. Nonetheless, these studies were the first to infer the presence of sites of frequent replication initiation in the germline, at S jumps, and the presence of lower efficiency origins within the N-domains. Subsequent studies mapping the timing of DNA replication throughout S phase showed that S-jumps also correspond with to sites of early replication timing in somatic cells, suggesting that many of these early, efficient replication zones are conserved across cell types (13, 14).

More recent approaches support the idea that replication initiates in kilobase-sized zones, but suggest that, within these zones, origin firing is spatially stochastic. Ok-seq maps Okazaki fragments synthesised on the lagging strand and provides a probability of being replicated as the lagging strand for all sites in the genome (15). High resolution repli-seq is based on a more elaborate replication timing analysis that takes advantage of a machine learning algorithm to identify sites where replication initiates (16). Both approaches provided signatures consistent with the firing of multiple low efficiency origins within defined initiation zones, rather than the imprint of individual highly efficient origins. This model is further supported by the recent application of optical replication mapping (ORM) (17).

However, the isolation and deep sequencing of short nascent leading strands (SNS-seq) identified ~ 300,000 discrete sites of DNA replication initiation (18, 19). As initiation sites were identified in increasing number and with better resolution, certain genetic and epigenetic features that correlated with origin firing were identified or confirmed, such as increased G/C content, marks of open chromatin and DNaseI hypersensitivity and histone H3K4 trimethylation (6, 18, 19). These experiments also led to renewed interest in the extent to which DNA sequence or DNA secondary structure, particularly G quadruplexes, are linked to, and may even define, sites of efficient human replication initiation (19–22).

The identification of individual sites of origin activity and the observation of broader zones of initiation are not, of course, contradictory but may rather reflect the resolution at which replication origin activation is studied. Nonetheless, the relationship between sites of efficient initiation, *i.e*. defined replication origins, and initiation zones remains unclear: do initiation sites contain discrete high efficiency origins or are they made up of a collective of stochastic low efficiency sites? Addressing this question requires a robust estimate of replication origin efficiency, *i.e*. the probability with which a replication origin fires during S-phase.

Here, we present a modification of initiation site sequencing (ini-seq) (23) that allows the high-resolution mapping of replication origins while simultaneously providing a quantitative estimate of initiation efficiency. Ini-seq is built on a cell-free system for the initiation of human DNA replication, in which human cell nuclei isolated from late G1 phase cells start DNA replication within a few minutes after the addition of a cytosolic extract from proliferating human cells *in vitro* (24, 25). In the original ini-seq method (23), nascent DNA was labelled by the addition of digoxigenin-dUTP, immunoprecipitated by an anti-digoxigenin antibody and sequenced alongside an input genomic DNA. This method allowed the identification of replication origins, but no assessment of origin efficiency was made. In the present study, we have added density substitution to ini-seq to separate replicated from unreplicated DNA, following the principles of the classical Meselson-Stahl experiment (26). We have used heavy bromo-dUTP (Br-dUTP) as a substitute for light dTTP during DNA replication *in vitro* and used density equilibrium gradient centrifugation to separate the replicated heavy/light DNA (HL) from unreplicated light/light DNA (LL). Following massive parallel sequencing of both HL and LL DNA fractions, and by the use of a custom-made algorithm, we identified 23,905 replication origins in nuclei of the human cell line EJ30. We show that these detected origins exhibit an excellent overlap with origins identified by SNS-seq in the same cell line, and also with an existing database of ‘core’ origins identified by SNS-seq in 19 other human cell types (27). There is also a good overlap with initiation zones determined by Ok-seq experiments performed in 9 different cell lines (15) (see also Material & Methods). Our combined origin localisation and efficiency information allows us to show that the most efficient origins in our set are not randomly distributed, but rather cluster at the edges of initiation zones, revealing a previously unappreciated fine structure to origin activity within these zones. We propose that these discrete, highly efficient origins explain much of the activity of the initiation zones and that these sites define constitutive sequences for replication initiation.

## Materials & Methods

### Cell culture, synchronisation and fractionation

Human EJ30 bladder carcinoma cells were cultured as monolayers and synchronized in late G1 phase by a 24-hour treatment with 0.6 mM mimosine (Sigma) as described (28). Template nuclei for DNA replication initiation *in vitro* were isolated from synchronized cells by hypotonic treatment, Dounce homogenisation and centrifugation as described (24, 28). Cytosolic S100 extract from proliferating human HeLa cells was purchased from Ipracell (Mons, Belgium).

### Initiation site sequencing (ini-seq)

The original protocol for ini-seq (23) was adapted to allow separation of replicated from unreplicated DNA by density substitution as follows:

### DNA replication reactions

Three identical reactions were run in parallel for each experimental condition. Each reaction contained HeLa cytosolic S100 extract (containing 400 μg of protein), 6 μl of template nuclei from late G1 phase EJ30 cells synchronised by mimosine (corresponding to 1.6-1.8 × 10^6^ nuclei), and a 5× premix of buffered nucleotides with an ATP regenerating system (yielding final concentrations of: 40 mM K–HEPES pH 7.8; 7 mM MgCl2; 3 mM ATP; 0.1 mM each of GTP, CTP, UTP, dATP, dGTP, dCTP, Br-dUTP; 0.5 mM DTT; 40 mM creatine phosphate; and 5 μg phosphocreatine kinase [all Merck]). The final volume of each reaction was adjusted to 100 μl with replication buffer (20 mM K-HEPES, pH7.8; 100 mM potassium acetate; 1 mM DDT; 0.5 mM EGTA). Reactions were incubated for 3 hr at 37 °C.

### Preparation of replicated DNA

DNA replication initiation reactions were stopped by dilution into 1 ml of ice-cold DNA buffer (10 mM Tris-Cl, pH8.0; 125 mM NaCl; 1 mM EDTA), and nuclei were pelleted at 16,000 × g for 5 min at 4°C. The pelleted nuclei were resuspended into 500 μl of DNA buffer and material from the three identical reactions was pooled at this stage. Nuclei were pelleted again, dissolved in 750 μl lysis buffer (10 mM Tris-Cl, pH8.0; 125 mM NaCl; 1 mM EDTA, 1% l-laurylsarcosine, 2mg/ml proteinase K) and incubated at 55 °C for 24 hr. High-molecular weight DNA was extracted with phenol/chloroform, precipitated with ethanol, and dissolved in DNA buffer at 55°C for 3 hrs. Concentrations were determined by spectrophotometry using a NanoDrop 1000 spectrophotometer (Thermo Fisher Scientific).

For DNA fragmentation, exactly 20 μg of DNA was adjusted to a volume of 130 μl per sample with DNA buffer and fragmented on a Covaris focussed ultrasonicator ME220 (using microTUBE-130 AFA Fiber Strips V2 and the following settings: 130 s duration, 70 W peak power, 20 % duty factor, 1,000 cycles per burst, average power of 14.0). Two samples were processed in parallel for each reaction, and subsequently pooled. Resulting distributions of fragmented double-stranded DNA were checked to the target size of 300 bp by neutral agarose gel electrophoresis.

### Equilibrium density gradient centrifugation

The pooled DNA preparations of each sample were adjusted to 4.9ml with 1.5 M Cs_2_SO_4_, 10 mM Tris-Cl pH 7.4, 1 mM EDTA (corresponding to a refractive index of 1.3700) and loaded into OptiSeal polypropylene centrifuge tubes (Beckman Coulter, 362185). Centrifugation was performed in a near-vertical NVT-90 rotor (Beckman Coulter) at 55,000 rpm at 20°C for 22 hours (no break) in a Beckman J6-MC ultracentrifuge (Beckman Coulter). Gradients were pumped out from bottom to top through a glass capillary tube attached to silicone tubing, using peristaltic pump P-1 (GE Healthcare) at a flow rate of 2 ml/min. Fractionation was performed manually at 6 drops per fraction. Refractive indexes of odd-numbered fractions were determined with an ATAGO R5000 hand refractometer. DNA concentrations of every fraction were determined by spectrophotometry using a NanoDrop 1000 spectrophotometer (Thermo Fisher Scientific). Samples were blanked against the caesium sulphate solution used for the gradients. Averages of at least four readings were taken for each fraction.

For each experiment two consecutive gradients were run to increase removal of contaminating LL DNA from HL DNA fractions. Raw DNA preparations were separated on a first density gradient. Fractions of the first gradient containing LL DNA (from the light side of the peak, RI = 1.3670 - 1.3660) were isolated and pooled. Fractions of the first gradient covering the HL peak area (RI = 1.3695 – 1.3715) were pooled and run again on a second density gradient. Fractions of the second gradient containing HL DNA (covering the peak and the adjacent fractions on either side, RI = 1.3705 - 1.3695) were isolated and pooled. DNA from the isolated pooled LL and HL fractions was desalted on PD MiniTrap G-25 columns (GE Healthcare) equilibrated in 10m Tris-Cl pH 7.4, 1 mM EDTA. Prior to library generation for DNA sequencing, the isolated LL and HL DNA fractions were concentrated by precipitation in 50% isopropanol, 0.5 M NH4-acetate, 2.5 μl/ml glycogen, washed in 70% ethanol, and dissolved in water.

### SNS-seq

Short nascent strand sequencing was performed as described in Akerman et al. (27) with the following modifications: After isolation of nascent strands by sucrose centrifugation, two sizes were recovered, a pool of 0.5 to 2 kb and a pool of 2 to 4 kb fractions. These were processed and sequenced separately.

### Library preparation & sequencing

Libraries were prepared with NEBNext^®^ Ultra™ II DNA Library Prep Kit for Illumina (NEB, E7645S) with 10 ng input DNA and 15 PCR cycles for amplification. Size selection was performed with an 8 % polyacrylamide gel stained with SYBR Gold (Thermo Fisher, S11494) with excision of fragments between 200 bp to 500 bp. DNA was extracted from the gel for 2 h at 37 °C in 0.5 M sodium acetate, 0.05 % SDS, precipitated in the presence of glycogen *v/v* with Isopropanol, washed once in ethanol 70 % and resuspended in water. Libraires were quantified with the KAPA Library Quantification Kit (Kapa Biosystems, KR0405) and sequenced on an Illumina HiSeq 4000 (50 bp single-end reads).

### Sequence alignment

Raw sequencing reads were aligned to the human genome assembly 38 (hg38) using `bowtie2` (29). Reads that were not uniquely aligned to one genomic position were removed. `Samtools` (30) was used to remove PCR duplicates and generate .bam and .bed files. Visual inspection of the data and screenshots were generated using IGV 2.9.4 (31).

### SNS-seq origin calling

SNS-seq origins in EJ30 cells were called with MACS2 using the parameters previously described by Akerman et al (27), with total genomic DNA from EJ30 cells used as input.

### Custom ini-seq 2 origin caller

We devised a custom script in R, calling subroutines in `bedtools` (32) and `Awk`, to call origins based on the reads in the HL and LL fractions and attribute to them an efficiency score. Briefly, we started by counting sequencing reads in 100 bp windows for both HL and LL fractions. Read counts were normalised to the total number of reads in each sequencing library. We then kept only those windows with a log_2_(HL/LL) read ratio ≥ 2. The remaining windows that were ≤ 500 bp from each other were merged. Only domains ≥ 200bp were retained. These domains we now call islands. We recounted the sequencing tags for all islands and normalised the counts to the total number of reads before calculating the efficiency score for each island (HL/(HL+LL)). For an island to be called an origin we set a cut off at ≥ 0.8 (that corresponds to a log_2_ ≥ 2). We finally computed the Z-score for each origin and used this to divide the origins into three equally sized groups, which we termed high, medium and low efficiency.

### Software and packages used

All scripts and statistical analyses were executed in R (33) or the Unix command line. Venn diagrams were created with the `bedtools closest` function and plotted with the venneuler package. For comparison of replication initiation sites between ini-seq 2 and other techniques, an intersect distance of 0 kb has been set for distributions of initiation sites larger than ini-seq 2 (Ok-seq, Bubble-seq and ORM). For comparison with SNS-seq and the original ini-seq experiment (23), whose distribution is within the same range as ini-seq 2, a maximum intersect distance of 5 kb was applied, and finally for the narrowly defined S-jump a distance of 10 kb was chosen. Ini-domains were generated using the `bedtools` merge function at a maximum distance of 100kb, only domains containing at least 6 origins were kept. Cluster analysis of high, medium and low efficiency origins was performed using the `clusterdist` option of `clusterscan` (34) with a distance for determining a cluster of 30 kb. Feature coverage around origins was plotted using `deepTools` (35).

### Venn diagram overlap statistics

Permutation analyses for testing Venn overlap significance were carried out using `regioneR` (36) with 10,000 permutations. At this number of permutations, the minimum *p* value is 10^−4^. We also report a Z-score that estimates the strength of the result (in this context, the Z-score is computed as the distance between the evaluation of the original region of interest and the mean of the random evaluations divided by the standard deviation of the random evaluations).

### Predictors of replication origin efficiency

Origin sequences were recovered from the hg38 human genome and base composition statistics were performed using the `Biostrings R/Bioconductor` package (37). Information about chromatin state at origins, *i.e*. DNaseI hypersensitivity and histone marks from H9 cells, data generated by the lab of Bing Ren (UCSD), was recovered from the ENCODE portal (https://www.encodeproject.org/, (38)). Non-B DNA-forming motifs, comprising direct repeats (DR), mirror repeats (MR), inverted repeats (IR), short tandem repeats (STR), Z-DNA (Z) and G-quadruplexes (GQ), were recovered from the non-B database (https://nonb-abcc.ncifcrf.gov/apps/site/default, (39)). Enrichment or depletion of each feature at replication origins was quantified by averaging the coverage signal over the origins obtained with the `computeMatrix` function of `deepTools` (35) using bin sizes of 10 nt and scaling origin regions to 5 kb. Reported correlations between predictors were computed as Pearson correlations using only origins with complete observations. The correlation heatmap was constructed by single-linkage clustering using Manhattan distances.

### Principal component analysis and statistical modelling

Principal component analysis (PCA) and the selection of predictive models were performed using the `caret` package in the R environment (40). We first assessed the correlations (using a threshold of |correlation| ≤ 0.85) and linear dependencies (using QR decomposition) in between the selected predictors and found that all selected predictors were independent. Predictor values were then centred and scaled, *i.e*. calculated as Z-scores, and the PCA was performed using the `prcomp` built-in command of R without consideration of efficiency values. Eigenvalues and vectors were visualised using the `fviz_pca_var` function of the `factoextra` R package. The 20 selected predictors were then used to build regression models predicting origin efficiency. To do this, predictor values were randomly assorted into two sets: 70% of the sets were used for training and the remaining 30% providing the testing set. The training set was used to select models using support vector machines with radial basis kernel function algorithm (`svmRadial` model from the `caret` package). Models were optimized by tuning the `svmRadial` parameters (sigma and C) over a 10-fold cross validation scheme. To assess their overall performance, the models were challenged against the test set and the model explaining the highest amount of variation on both the training and test sets was selected.

### External data sources & processing

S-jump data were taken from Huvet et al (12) and lifted over from hg19 to hg38 keeping all N domains where the size change was < 5%. The position of the S-jump was arbitrarily mapped at 1 nucleotide resolution to the edges of the N-domains. This approach identified 1018 S-jumps. Bubble-seq data were taken from Mesner et al (41) and ORM data from Wang et al (17). Replication timing data were taken from Rivera-Mulia et al (42). SNS-seq data were taken from Akerman et al (27), using the existing categorisation of ‘core’ and ‘stochastic’ origins proposed by the authors. Finally, Ok-seq data were retrieved from https://github.com/CL-CHEN-Lab/OK-Seq (15, 43), from which we used data from the nine cell lines BL79, IARC385, K562, IB118, IMR90, Raji, TLSE19, GM0699, and HeLa. For these data sets, we defined core initiation zones as being present in eight out of the nine cell lines. The remaining zones were classified as stochastic.

## Results

### Ini-seq 2

In order to study the localisation and efficiency of DNA replication origins, we adapted the initiation site sequencing (ini-seq) method (23). Ini-seq is based on the ability of human cell nuclei to initiate DNA replication *in vitro* upon the addition of a cytosolic extract from proliferating human cells (24). Under these experimental conditions, DNA replication is initiated, and replication forks move away from replication origins with much reduced fork speeds than observed *in vivo* (44). In our new modified approach, we chose to label nascent DNA by incorporation of heavy bromo-dUTP (Br-dUTP) as a substitute for the light dTTP, rendering semi-conservatively replicated DNA heavy-light (HL) and leaving unreplicated DNA light-light (LL). We thus isolated late G1 phase nuclei from mimosine-arrested human EJ30 cells, and initiated DNA replication *in vitro* in the presence of Br-dUTP instead of dTTP (Figure 1A). Following isolation of total DNA and sonication, the LL and HL DNA fragments were separated by caesium sulphate gradients (Figure 1B), using parameters previously established (45, 46).

**Figure 1.**
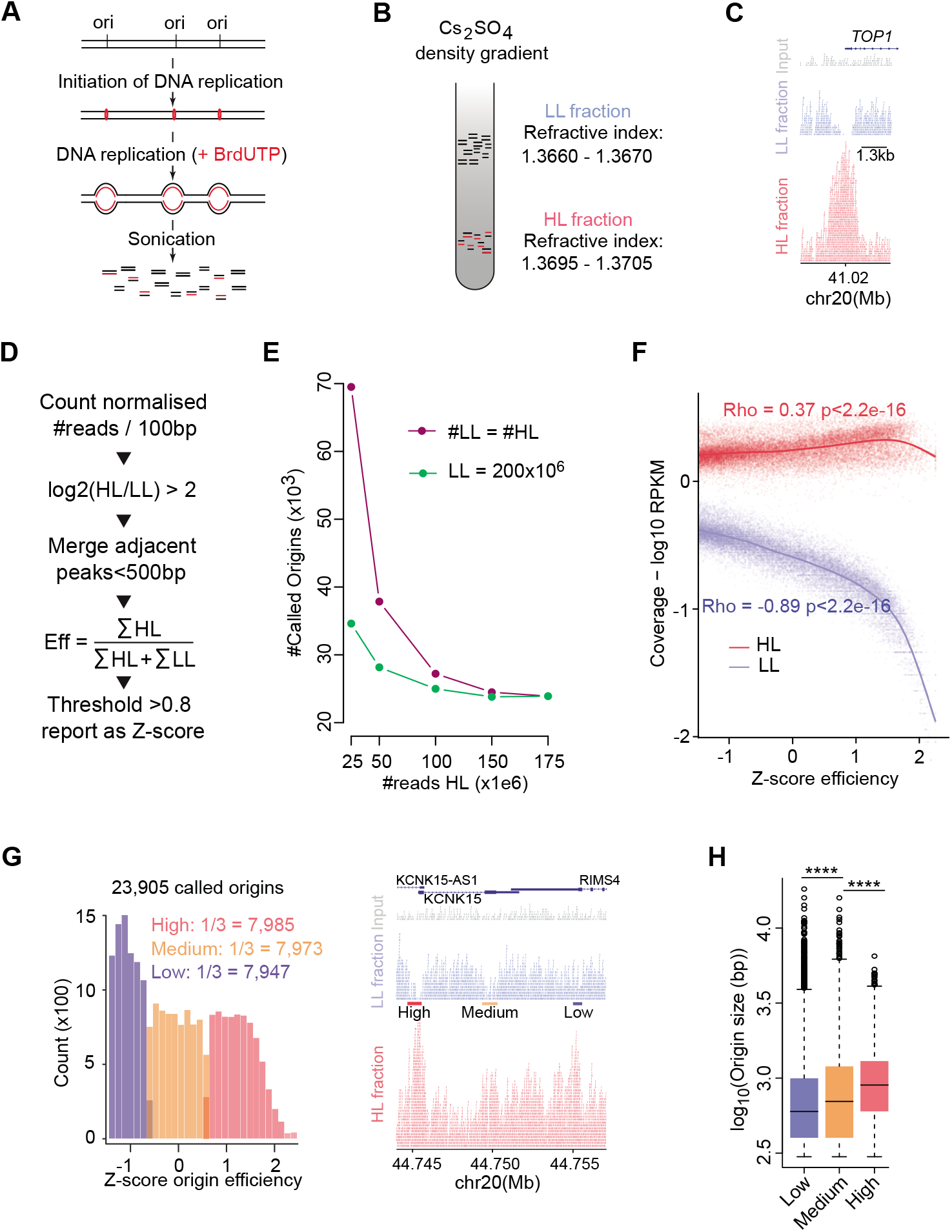
Ini-seq 2: a method allowing fine mapping of replication origins and their efficiency. A. Schematic representation of the ini-seq 2 workflow to label active DNA replication origins in nuclei from EJ30 cells by density substitution. B. Separation of the newly synthesised (heavy/light, HL) and unreplicated (light/light, LL) DNA by equilibrium density centrifugation in caesium sulphate gradients. C. An IGV screenshot demonstrating enrichment of HL reads (red) and depletion of LL reads (blue) reads at a well-studied origin near the TSS of the *TOP1* gene. Total input DNA is represented in grey. D. Diagrammatic representation of the custom algorithm developed to call ini-seq 2 replication origins. E. Number of called origins as a function of the number of reads sequenced. Purple line: reads in HL = reads in LL; green line: a fixed number of LL reads (200 × 10^6^) while the number of HL reads is varied. F. Read coverage for each origin as a function of efficiency. Red = HL, blue = LL. Correlation: Pearson. G. Classing of origin efficiencies. (Left) Distribution of origins by their efficiencies and binning of equal numbers into high, medium and low classes. (Right) IGV screenshot showing a genomic region containing examples of the three classes of origins. HL reads are shown in red; LL reads in blue and total input DNA reads in grey. H. Distribution of origin size by class. **** = p < 2.2 × 10^-16^; K-S test.

Following massive parallel sequencing and read alignment of LL and HL DNA fractions, clear peaks were observed in the HL fraction at sites of replication origins, exemplified by the well-documented origin at the transcription start site (TSS) of the *TOP1* gene (Figure 1C; 47). Interestingly, we also observed a depletion of LL reads at the centre of many HL peaks, with no reads at all at some loci, as at the *TOP1* origin (Figure 1C). This observation implies that the vast majority of nuclei had initiated replication at these sites and that most if not all of the parental LL DNA was converted to replicated HL DNA. Importantly, this substantial local conversion of LL to HL DNA suggests that the LL fraction cannot be treated as a conventional ‘input’ for computational normalisation of the HL signal, and for peak calling. We therefore created a custom origin calling algorithm that exploits the concomitant increase of the HL and depletion of LL reads to compute not only the location of each origin, but also an efficiency score of origin activation for that site (Figure 1D). Biological duplicates of this experiment exhibited both a high overlap of locations and a positive correlation in efficiency scores for these origins (Figure S1A & B). Therefore, we pooled the data from both replicates to yield 175 × 10^6^ HL reads and 200 × 10^6^ LL reads and identified 23,905 unique origins. We then compared this dataset to the origins that were called using MACS2 (48), a conventional peak caller designed for the analysis of ChIP-seq data. When applied to the HL fraction using the parameters of Akerman et al (27) (Figure S1C), this analysis showed that practically all the origins identified by our custom peak caller were a subset of those called by MACS2. Further, we find that using the LL fraction in our peak-caller improves the dynamic range of the efficiency scores (Figure S1D). We also compared our data with the original ini-seq dataset (23), and observed 75% overlap (Figure S1E).

Our custom peak caller did not call any further origins beyond a read depth of ~ 150 × 10^6^ for both HL and LL (Figure 1E). Notably, by fixing the number of LL reads at 200 × 10^6^, only ~ 50 million HL reads were needed to reach saturation (Figure 1E), suggesting that depletion of LL reads is important in calling origins. We further explored this idea by examining read coverage as a function of origin efficiency (Figure 1F), which highlighted a much greater correlation with LL than HL reads. Together, these observations highlight the importance of sequencing both replicated HL and unreplicated LL DNA for determining origin efficiency. Additionally, this approach avoids potential biases introduced by sequencing a traditional ‘input’ sample (49), as the number of reads in HL and LL samples are inter-dependent (Figure 1F). We split the set of 23,905 origins determined by our custom peak caller into three equally-sized groups, which we named according to their relative efficiency as ‘high’, ‘medium’ and ‘low’ (Figure 1G). We observed that the size of an origin increases with its efficiency class (Figure 1H), with an overall median size of 699 bp. As identification of high efficiency origins is mostly driven by the local depletion of LL read coverage (Figure 1F), the positive correlation of origin efficiency with size argues that the observed localised depletion of LL DNA at these sites is unlikely to occur by chance. Further, the distribution of origin size is well within the range expected for highly defined isolated origins rather than representing a large domain of multiple origins. In conclusion, this new density substitution-based ini-seq 2 method provides information both on origin location and efficiency.

### Ini-seq 2 origins are a subset of origins identified by SNS-seq in EJ30

In order to benchmark ini-seq 2 against an established DNA replication origin identification method performed on the same cell line, we carried out short nascent strand sequencing (SNS-seq) on EJ30 cells. We isolated nascent strands in two fractions (0.5 – 2 kb and 2 – 4 kb) by gel electrophoresis (50) and individually sequenced them to provide additional confidence for peak calling. An example of mapped raw reads is shown in Figure S2. Using the pipeline established by Akerman et al (27), we identified 175,536 peaks, which included 88% of the origins mapped by ini-seq 2 (Figure 2A). Furthermore, we found that the origins called using the ini-seq 2 pipeline were more tightly defined that those of SNS-seq, at two different SNS DNA fraction sizes, as seen in the normalised coverage of reads around each called origin (Figure 2B) and by the size distribution of origins called by each method (Figure 2C). We conclude that the vast majority of origins identified by ini-seq 2 are confirmed by SNS-seq in the same cell line.

**Figure 2.**
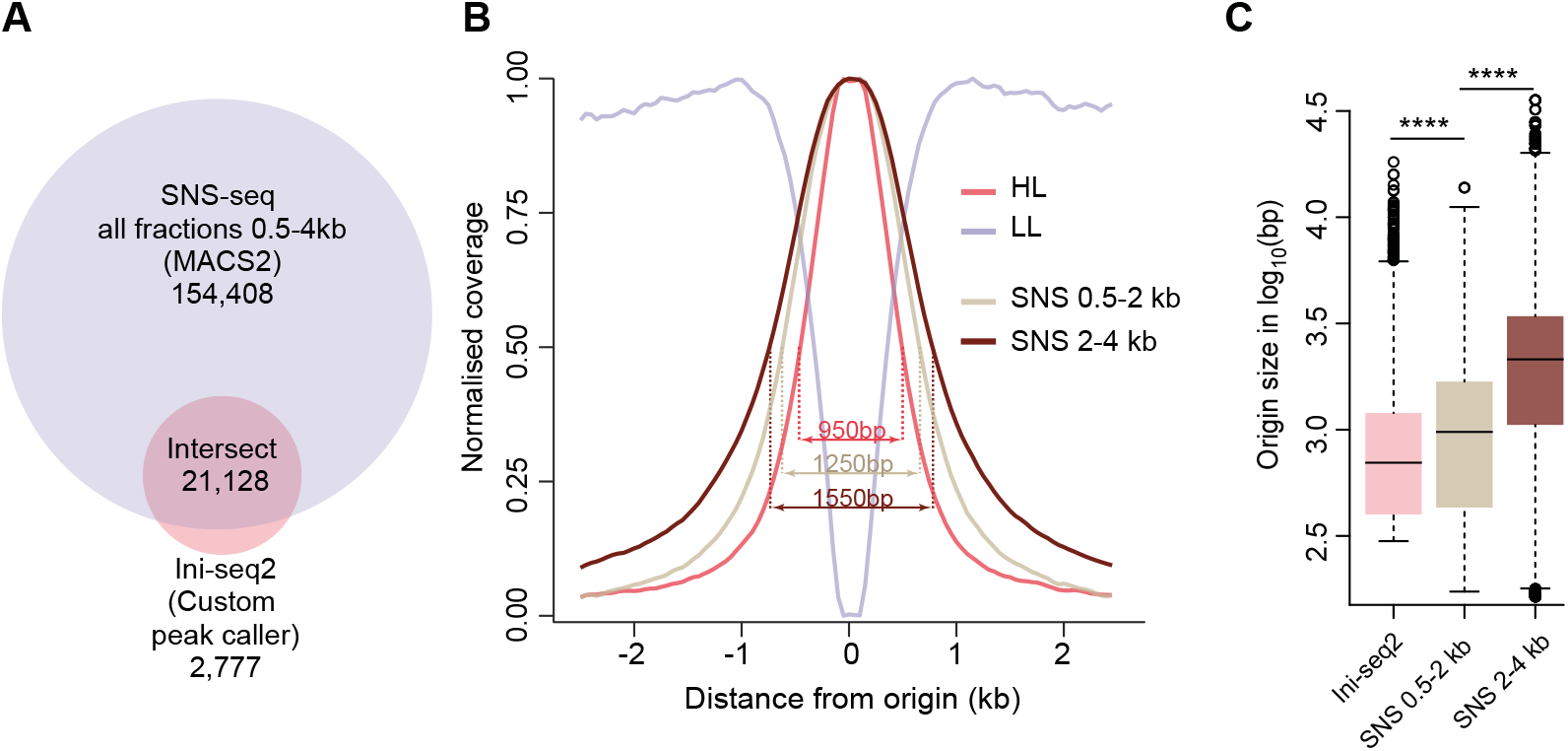
Comparison of ini-seq 2 and SNS-seq in EJ30 cells. A. Overlap of origins called by SNS-seq and ini-seq 2. SNS-seq was performed on two fractions, 0.5 – 2kb & 2 – 4kb, which were pooled. Permutation test p = 0.0001, Z-score 538. B. Read coverage around ini-seq 2 and SNS-seq origins. Width values are for the half height of each distribution. C. Distribution of origin size defined by ini-seq 2 and SNS-seq. **** = p < 2.2 × 10^−16^; K-S test.

### Ini-seq 2 identifies ubiquitous and constitutive early origins

We next compared the set of origins mapped in EJ30 cell nuclei using the ini-seq 2 pipeline with previously reported origin maps of the human genome derived from other cell lines. We first compared the size of replication initiation sites determined by each published technique (Figure S3A), in order to set an allowable overlap distance to compute the intersect distances between the sites mapped by each technique and ini-seq 2 (see Materials & Methods). We further validated our maximal cut-off distances by determining the dependence of the extent of intersect on the maximum intersect distances (Figure S3B). Thus, among the 1,018 S-jumps we could map to the hg38 genome assembly (see Material & Methods), 74% overlapped with ini-seq 2 origins (Figure 3A & B). We then compared our ini-seq 2 dataset with both bubble-seq (41) and optical replication mapping (17) and observed a modest overlap of 44% and 26%, respectively (Figure S3C & D). A recent study applying SNS-seq to 19 human cells lines (27) allowed the identification of a subset of origins, termed ‘core’ origins, that are present in all cell lines suggesting that they are highly conserved across tissues. In contrast, ‘stochastic’ origins are found in only one or a few of the cell lines. While 74% of the ini-seq 2 origins were found within the SNS-core origin group, only 19% were found in the SNS-stochastic group, leaving 7% in neither group (Figure 3C). Finally, we compared the ini-seq 2 dataset with published Ok-seq data. Applying the same classification approach as used by Akerman et al (27) to the published Ok-seq data from 9 different cell types (15, 43 and see Materials & Methods), we identified 2,103 ‘core’ Ok-Seq initiation zones of which 80% include ini-seq 2 origins. In contrast, of the 19,576 identified ‘stochastic’ Ok-Seq zones, only 25% include an ini-seq 2.0 origin (Figure 3D). Taken together these observations show that origins defined by ini-seq represent a subset of narrowly defined origins that overlap with core origins and cover core replication zones as identified by independent methods and across multiple cell types.

**Figure 3.**
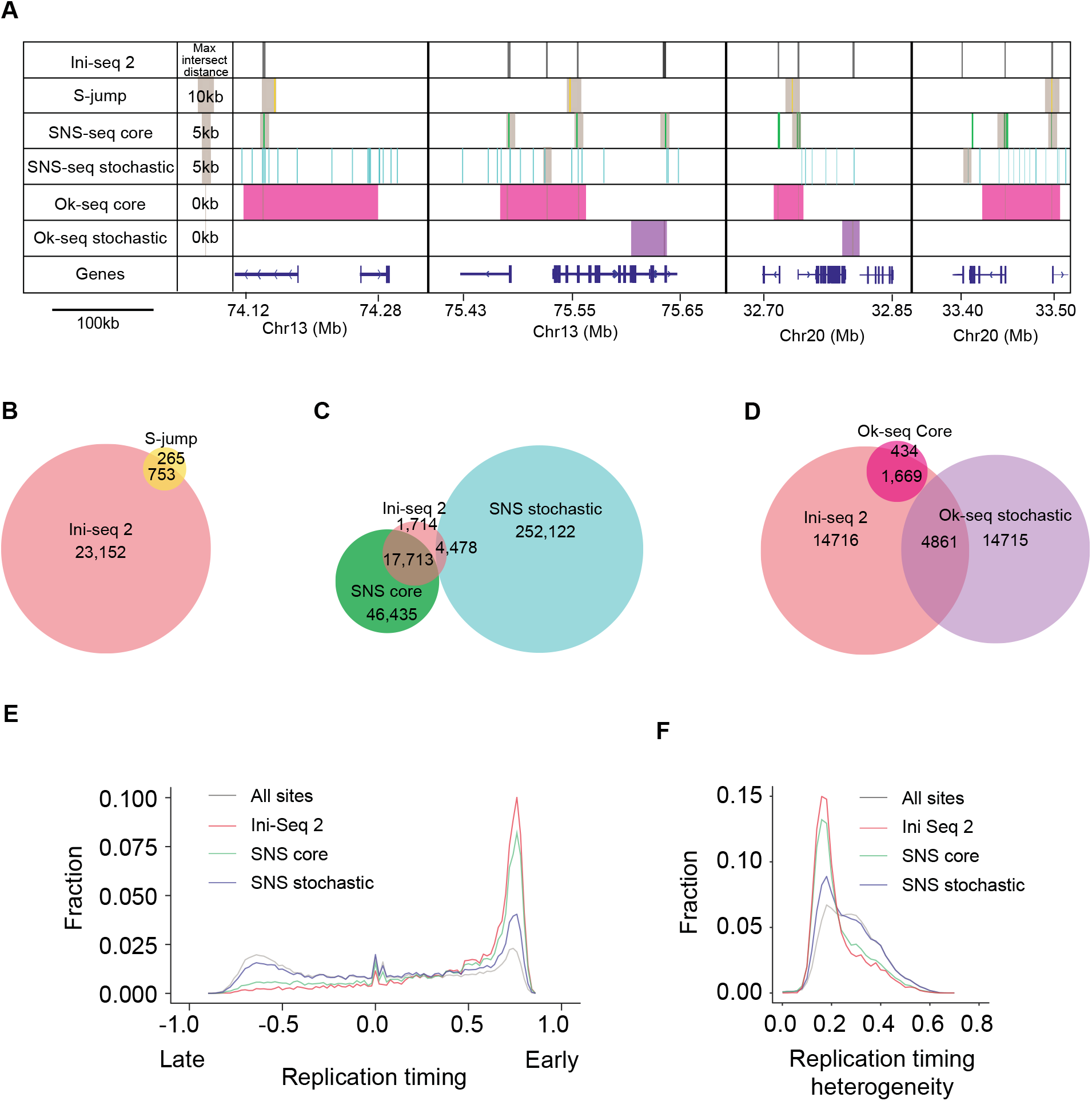
Comparison of ini-seq 2 origins mapped in other cell lines. A. Four representative genomic regions illustrating the position of origins identified by ini-seq 2, S-jumps, SNS-seq and Ok-seq. Grey boxes indicate the maximum distance that has been allowed to accept an intersect, computed based on the average size of the origins called by each method (see Materials & Methods). B - D. Venn diagrams showing the overlap between (B) ini-seq 2 origins and S-jumps, permutation test p = 0.0001, Z-score 58; (C) ini-seq 2 origins and SNS-seq, permutation test for core and stochastic respectively p = 0.0001, Z-score 379 and p = 0.0001, Z-score 217; (D) ini-seq 2 origins and Ok-seq, permutation test for core and stochastic respectively p = 0.0001, Z-score 89 and p = 0.0001, Z-score 61. E. Distribution of origins determined by ini-seq 2 and SNS-seq as a function of replication timing. F. Distribution of origins determined by ini-seq 2 and SNS-seq as a function of replication timing heterogeneity observed across nine cell lines (see Materials & Methods).

Using previously reported replication timing data average from nine cell lines (51 and see Materials & Methods) we found that both the ini-seq 2 and SNS-core, and, to a lesser extent the SNS-stochastic origins, are enriched in early replicating regions (Figure 3E). They are additionally characterised by low replication timing heterogeneity (*i.e*. conservation of replication timing between cell types (51 and see Materials & Methods)), in contrast to the stochastic origins (Figure 3F). Altogether, we conclude that origins identified by ini-seq 2 constitute a group of early replicating origins, active independently of cell line and type, and their early timing is not simply a feature of the origins identified by the ini-seq 2 technique.

### Determinants of origin efficiency

We next examined whether the efficiency information provided by ini-seq 2 could be used to refine our understanding of genomic and epigenetic features that help determine origin efficiency. We first examined the correlation between base composition and origin efficiency. We observed a positive correlation between the GC content at the origin and their efficiency, with GC contents of 0.63, 0.65 and 0.75 for low, medium and high origins respectively (Figure 4A), and, consequently, we also observed a correlation with CpG islands (Figure S4A). Furthermore, we note that highly efficient origins are found within AT-rich regions (0.46 GC / 0.54 AT content), a feature that is not observed at domains displaying medium (0.49 GC / 0.51 AT content) and low (0.53 GC / 0.47 AT content) origins (Figure 4A).

**Figure 4.**
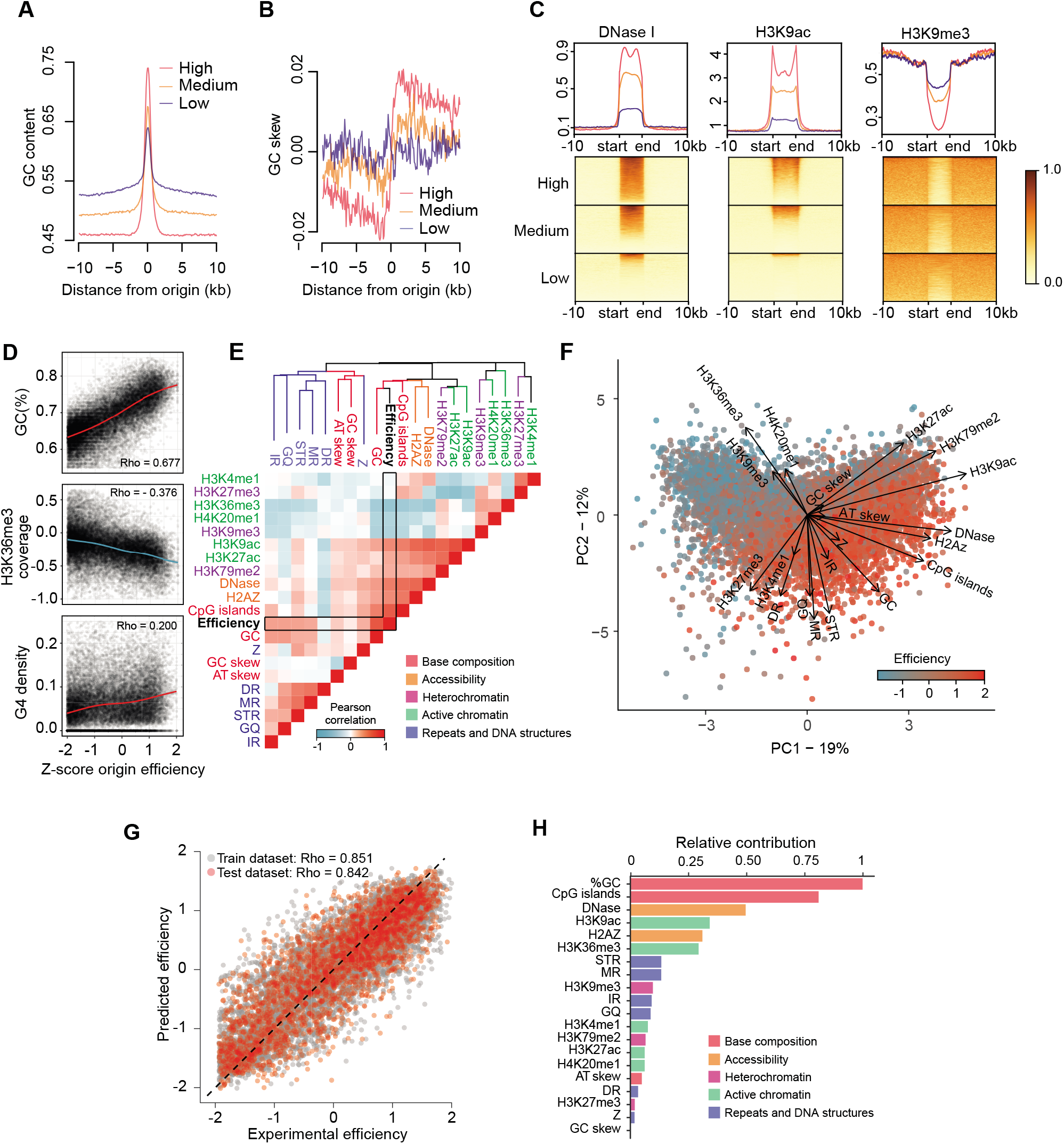
Determinants of origin efficiency. A. GC content in a 20kb window around origin centres for the three efficiency classes of ini-seq 2 origins. B. GC skew (G − C / G + C) computed in 100 bp bins in a 20kb window around the origin centre for the three efficiency classes of ini-seq 2 origins. C. Coverage plots for DNase I hypersensitivity, H3K9 acetylation and H3K9 trimethylation within and 10 kb around the three efficiency classes of ini-seq 2 origins. Origin lengths were scaled and are defined by ‘start’ and ‘end’ labels. D. GC content, H3K36 trimethylation and G4 density as a function of origin efficiency. Correlation: Pearson. E. Heatmap reporting the correlation between pairwise combinations of origin features. Blue = negative Pearson correlations; Red = positive Pearson correlations. The dendrogram is generated using an unsupervised clustering algorithm based on distances computed from Pearson correlations (see Material & Methods). The colours of the branches denote the five types of origin features. F. Principal component analysis of origin efficiency using features described in panel (E), highlighting the strength and direction, *i.e*. eigenvectors, for the contribution of each feature to origin efficiency. G. A statistical model allows prediction of origin efficiency using these features as predictors. Origins used to train and test the model are depicted in grey and red, respectively. E. Quantitative estimate of predictor contribution to the statistical model. The colours of the bars denote the five types of origin features.

We then examined the nucleotide skew around origins in the three efficiency classes. We observed that the ‘high’ efficiency origins globally exhibit a marked transition in GC and AT skew (Figure 4B & S4B). This skew transition is less evident in the medium and absent in the low efficiency group, suggesting that the highly efficient origins are drivers for the observed skews. The much-reduced skews score in the low efficiency group suggests that they are indeed less likely to fire in each cell cycle.

Since non-B-form DNA has been proposed to be linked to the presence of origins (19, 22, 52, 53), we assessed the impact of the presence of repetitive and structure-forming DNA sequences at and around origins in the three efficiency classes (Figure S4B). We found an efficiency-dependent enrichment of G4s, inverted repeats, mirror repeats and short tandem repeats at origins, suggesting secondary structure formation may promote origin activity, possibly by creating a more accessible chromatin environment. Taking datasets from the ENCODE project (38 and see Materials & Methods), we find that, consistent with this idea, the most efficient origins have the greatest DNaseI accessibility, the signal for which is very sharply defined at our called origins and correlated with origin efficiency (Figure 4C). Further, histone modifications associated with accessible chromatin, such as H3K9ac, are also enriched, while those associated with heterochromatin, such as H3K9me3, are excluded (Figure 4C & S4C).

We next examined the more general correlation of local genetic and epigenetic features at and around DNA replication origins and the efficiency with which they fire. We first assessed the correlation of 20 genetic and epigenetic features of replication origins with firing efficiency. We found both positive and negative linear correlations, as illustrated by the correlation with the GC content and H3K9me3 coverage, respectively (Figure 4D). There are also more complex correlations. For example, while a moderate positive correlation is found between G4 density and origin efficiency, it is clear that some origins that do not contain G4-forming sequences can be very efficient (Figure 4D). We then calculated the correlation coefficient for all pairwise combinations of features, including origin efficiency, and clustered the features using these correlation coefficients as a distance metric. Surprisingly, we found that features of similar classes (*i.e*. base composition, DNA accessibility, active/inactive chromatin marks, and DNA structures) cluster together (Figure 4E), suggesting that a single feature cannot explain the wide range of observed origin efficiency and that rather a combination of features is more likely to determine origin efficiency. We used principal component analysis (PCA) to investigate further the inter-dependencies among the different origin features (Figure 4F). This revealed that the contribution of these five classes of features can be orthogonal within the plan of the PCA, *i.e*. their contributions are independent rather than additive. For example, the contribution of DNA structures, such as G4s, is orthogonal to the active histone marks, such as H3K9ac, suggesting that the absence of DNA structures at an origin can be compensated by an active chromatin context for an origin to be efficient.

We finally aimed to quantify the contribution of each feature to origin efficiency. To do this, we devised an unsupervised machine learning algorithm to predict origin efficiency based on the set of these 20 predictors. We constructed regression models using 70 % of the origins as training sets with the remainder acting as testing sets. We selected a model which allows prediction of the efficiency of both the training and testing origin sets with high accuracy (Pearson correlation *Rho* ~ 0.85, Figure 4G), which is comparable to the correlation between two biological replicates of ini-seq 2 (Pearson correlation *Rho* ~ 0.77, Figure S1A). Our model hence shows that this set of features is an efficient predictor of origin efficiency. We observed that the key determinants of origin efficiency are base composition and accessible chromatin, but we note that DNA secondary structures collectively make a significant contribution with no individual structural class standing out (Figure 4H).

### Ini-seq 2 reveals a higher order organisation of replication origins by efficiency

We next assessed the genome-wide distribution and organisation of ini-seq 2 origins according to their efficiencies. We started by comparing their distribution within N-domains and found that the density of the high efficiency class origins is increased at N-domain borders, whereas the medium and low efficiency classes are found to be weakly, or not enriched at these sites, respectively (Figure 5A). We then asked if a similar organisation is seen in the much smaller domains defined by Ok-seq initiation zones (Figure S5). We found that the high efficiency origins are closely localised to the border of the Ok-seq initiation zones, that the medium efficiency origins are distributed within the zones while the low efficiency origins are equally distributed within and outside the zones (Figure 5B). We then asked whether we could define origin-rich domains *ab initio* using our ini-seq 2 data by simply merging all origins within 100 kb of each other, regardless of their efficiency class (see Material & Methods). This exercise generated 699 domains, which have a median size of 205 kb (Figure S5) and included 15,942 of the total 23,905 ini-seq 2 origins. Then we asked how the origins are organised by efficiency class within those domains and found that the high efficiency origins are also preferentially enriched, compared to the medium and low efficiency origins, at their borders (Figure 5C & D). We also observed a reduction of high efficiency origins within the domains (Figure 5E). This implied that the highly efficient origins tend to be isolated by at least 100kb on either their 5’ or 3’ side. To test this hypothesis, we computed the inter-origin distance by efficiency class and found that the most highly efficient origins are significantly further apart from each other (median distance 130 kb) than the medium (median distance 68 kb) or low efficiency origins (median distance 25 kb) (Figure 5F).

**Figure 5.**
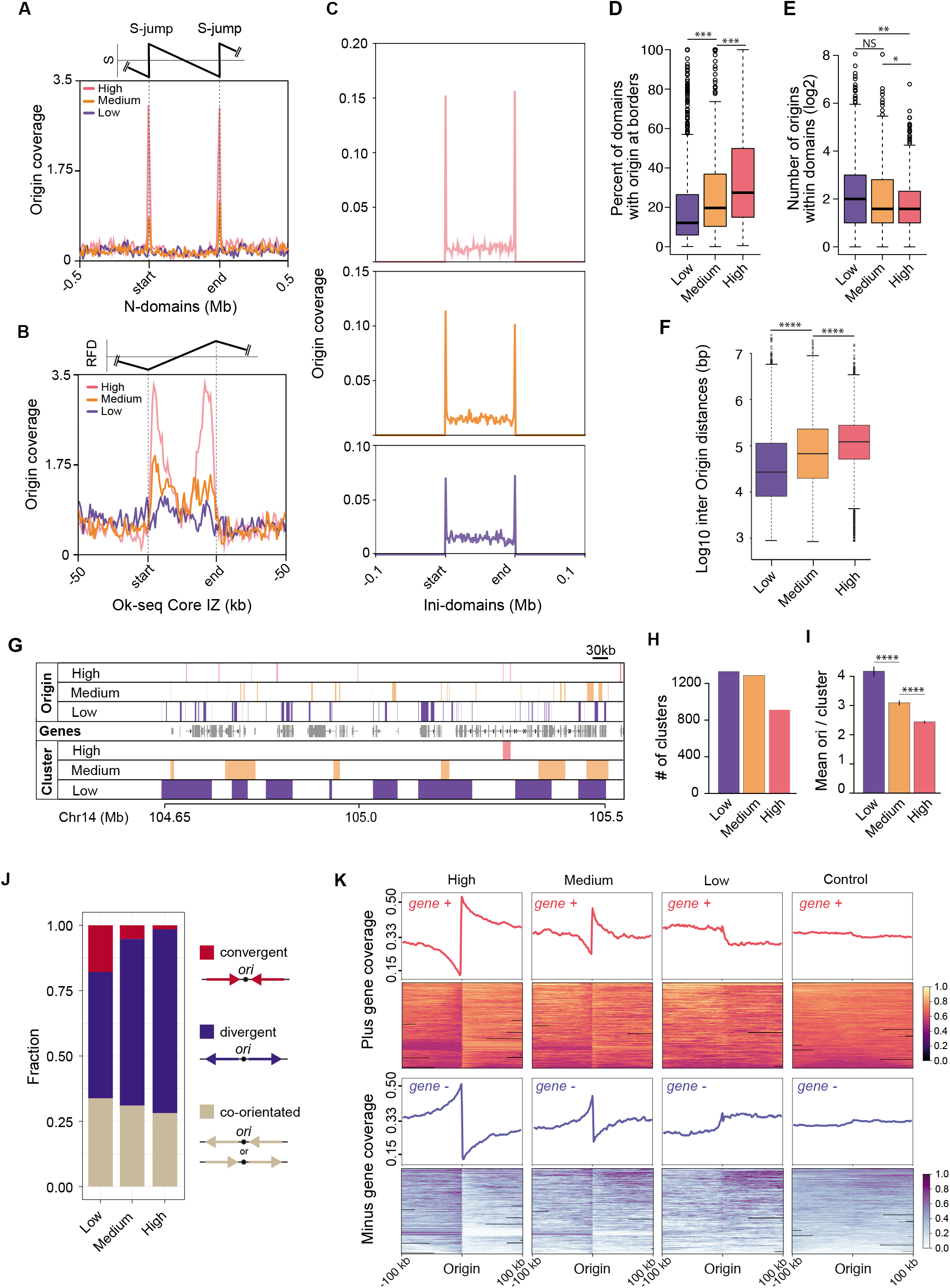
The higher order organisation of replication origins by efficiency. A. Origin coverage grouped by efficiency in normalised N-domains +/− 500 kb. B. Origin coverage grouped by efficiency in Ok-seq core initiation zones +/− 50 kb. C. Origin coverage grouped by efficiency in Ini-domains +/− 100 kb. D. Percentage of ini-domains with origins of each efficiency class at their boundaries. *** = p < 1 × 10^-7^; K-S test. E. Number of origins of each efficiency class within the ini-domains (borders analysed in D. were excluded). ** = p < 1 × 10^-5^; * = p < 1 × 10^-3^; K-S test. F. Inter-origin distances grouped by origin efficiency class. Central bar = median; whiskers = interquartile range. **** = p < 2.2 × 10^-16^; K-S test. G. Origin clustering. Example IGV screenshot showing the clustering of ini-seq 2 origins by efficiency in a ~ 1Mbp region of Chromosome 14. Top three lanes: mapping of the three efficiency classes of ini-seq 2 origins. Middle lane: genes. Lower three lanes: Origin clusters determined using the clusterdist function of clusterscan (34), set at 30 kb (see Materials & Methods). H. Number of clusters found in each ini-seq 2 origin efficiency class. I. Mean number of origins per cluster for each efficiency class (**** = p < 5 × 10^-12^ for low vs medium; p < 7 × 10^-15^ for medium vs high. J. Quantification of the orientation of the first gene either side of an origin, grouped by origin efficiency class. Gene orientation of the two adjacent genes is classed by the direction of transcription as convergent, divergent or co-orientated. K. Gene orientation coverage around the three classes of ini-seq 2 origins and a randomised control compared with 8000 randomly picked positions from a pool of genomic locations that are equidistant from two origins.

Finally, we examined the behaviour of ini-seq 2-defined origins at a smaller clustering range of 30 kb, comparable to Ok-seq initiation zones. By using a clustering algorithm (Figure 5G), we found that the low and medium efficiency origins form more clusters than those of high efficiency (Figure 5H). Consistent with this observation, we found that when clusters of highly efficient origins are found they contain fewer origins than the low and medium class (Figure 5I). Thus, the organisation of origins we have uncovered, in which high efficiency origins are found at initiation domain borders, is independent of the scale and type of the domains examined.

### Gene orientation is organised around the most efficient origins

Huvet and colleagues previously proposed an organisation of transcriptional units around origins defined by the borders of N-domains, that would avoid head-on collisions between transcription and replication (12). We thus wondered whether our more numerous high efficiency replication origins exhibited a similar behaviour. We therefore examined the orientation of the transcriptional units flanking the ini-seq 2 origins of each efficiency class. We quantified the proportions of convergent, divergent or unidirectional orientation of the first transcriptional units flanking the ini-seq 2 origins for each efficiency class (Figure 5J). Then we analysed more broadly the orientation of transcriptional units within 200 kb domains centred on the origins in each of the three efficiency classes (Figure 5K). These analyses revealed a strong preference for divergent orientation both for gene pairs and transcriptional units from replication origins around the high and, to a lesser extent, the medium efficiency class origins. This feature that is not shared with the low efficiency class origins, or when origin position is reshuffled (see Material & Methods). These observations show that the highly efficient origins define constitutive domains for the organisation of the human genome that tends to optimise the activities of DNA replication with transcription to favour co-directionality of replication fork and transcription.

## Discussion

The elusive nature of replication origins can perhaps be best exemplified by the ~100 kb initiation zone of the Chinese hamster dihydrofolate reductase locus, where decades of work led Joyce Hamlin to suggest a model ‘… *in which the mammalian genome is dotted with a hierarchy of degenerate, redundant, and inefficient replicators at intervals of a kilobase or less, some of which may have evolved to be highly circumscribed and efficient*.’ (3). In the present study we show that this hierarchical concept applies across the human genome (Figure 6).

**Figure 6.**
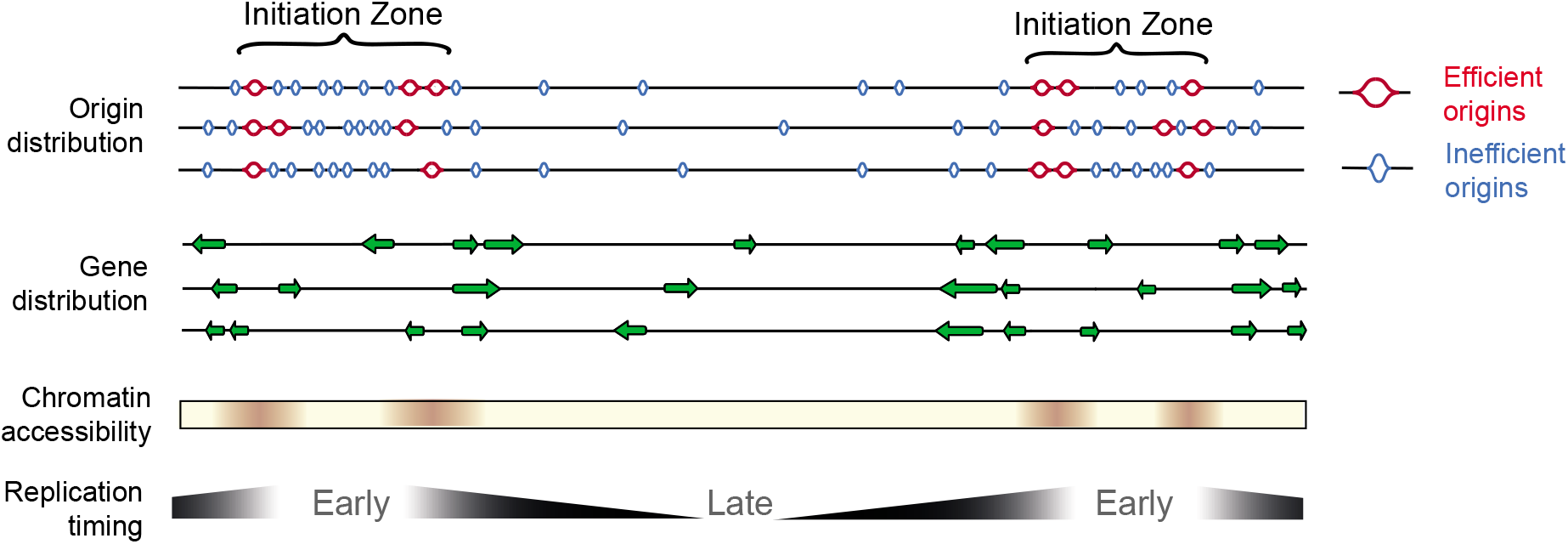
Model for the organisation of replication initiation zones in the human genome. Ini-seq 2 origins define early replicating regions of the genome with an ‘open’ chromatin structure. The borders of initiation zones are defined by the most efficient origins and local gene organisation that will minimise head-on transcription *i.e*. the core of the initiation zone is depleted in genes but enriched at the boundary with genes co-orientated with the direction of leading strand replication.

The improved version of ini-seq (23) we report here provides a high-resolution map of replication origins in a human cell line with direct quantitation of their activation efficiency. Our approach monitors the proportion of sequence reads from a given genomic location that have or have not taken up Br-dUTP as a consequence of semi-conservative DNA replication following initiation *in vitro*. The proportion of replicated to unreplicated DNA at a given locus therefore provides a direct readout of the efficiency by which the locus is replicated under these experimental conditions. Surprisingly, we not only found that DNA replication origins are marked by an enrichment of reads in the HL fraction, but for some origins the unreplicated LL fraction was completely depleted of reads at the same site (see *e.g*. the *TOP1* origin in Figure 1C). These areas of LL read depletion are comparatively short, despite the three-hour duration of the *in vitro* replication reaction, consistent with replication fork escape from the immediate initiation site being comparatively non-processive in this system.

By implication, our ini-seq origin efficiency reported here also provide a readout of the fraction of nuclei that have replicated their DNA at a given locus under the condition of the experiment. With the caveat that very small quantities of unreplicated DNA may have escaped detection by sequencing, the observation of unreplicated DNA depletion at efficient origins suggests that (almost) all nuclei must have initiated DNA replication at these locations. When analysed by immunofluorescence microscopy, only about 40-70% of template late G1 phase nuclei usually have established replication foci under these experimental conditions *in vitro* (24, 25, 44). The higher per-nucleus efficiency of initiation at the subset of highly efficient origins found here by ini-seq 2 is not inconsistent with these published microscopy data because the very short segments of replicated DNA at some origins monitored by ini-seq would fall short of the detection threshold of effective replication foci by immunofluorescence microscopy. In other words, nuclei that are not able to incorporate sufficient modified nucleotide for detection by immunofluorescence microscopy are nonetheless able to initiate at the most efficient origins.

Importantly, the highly efficient origins detected by ini-seq 2 form a subset of the initiation sites defined by the core origins previously detected by SNS-seq, and defined as core origins, across nineteen human cell lines (27), and by OK-seq across nine cell lines (15, 43). We have thus uncovered a class of replication origins, which we propose represent the most efficient (*i.e*. most likely to be used) sites of replication initiation in the human genome. Furthermore, these initiation sites can be delineated to a size of just a few hundred base pairs because of the resolution of ini-seq 2 and SNS-seq. Both techniques generate an average object size of some two orders of magnitude smaller than those of alternative techniques such as OK-Seq (15, 43) or optical replication mapping (17) (Figure S3A). Our observations therefore support the presence of highly efficient and tightly localised sites of replication initiation within the human genome.

We have also gained more insight into the genetic and epigenetic features that contribute to origin activity (reviewed in 54) by showing the degree to which each contributes to efficiency. Our ability to use the correlations between efficiency and genetic / epigenetic features around origin sites allowed us to train a machine learning algorithm to predict origin efficiency to an accuracy comparable to the experimental noise of the ini-seq 2 method (compare Figures S1B & 4G). We observe that, collectively, broader DNA secondary structure forming potential can contribute almost as much to driving origin efficiency as GC content and chromatin accessibility (Figure 4H). Indeed, there may be several routes to create an efficient human origin. Supporting this idea, structure forming potential largely clusters independently of GC content and ‘active’ chromatin marks (Figure 4E). This further suggests that, collectively, these features are sufficient to explain origin efficiency but that no one feature alone is sufficient to specify a site of efficient replication initiation.

Finally, we have established a size-independent principle for the organisation of origins into initiation zones that applies from Ok-seq domains at ~ 30 kb to N-domains at ~ 1 Mb. It is tempting to speculate that the subset of high efficiency origins play a major role in defining these zones and dictate the prevailing direction of replication out of them. Thus, this higher order organisation helps unify the concept of initiation zones with the notion of discrete replication origins in human cells (Figure 6).

## Data availability

Sequencing data can be accessed at the Gene Expression Omnibus archive with the accession number GSE186675. The code for the ini-seq 2 origin caller can be found at https://github.com/Sale-lab.

## Funding

Work in the Sale group is supported by a core grant to the LMB by the MRC (U105178808). H.S.W. received an award from the Annual Bursary Fund at St Catherine’s College Cambridge. L.K.L. received a 1-year Science Without Borders postdoctoral funding from the Coordenação de Aperfeiçoamento de Pessoal de Nível Superior (CAPES, Brasília, DF, Brazil). Work in the Krude group is supported by the Department of Zoology, University of Cambridge.

## Acknowledgements

We would like to thank Toby Darling and Jake Grimmet in Scientific Computing at LMB for support, Christelle de Renty, Jo Yeeles, Madan Babu and members of the Sale lab for discussions and comments on the manuscript.

## SUPPLEMENTARY INFORMATION FOR

**Figure S1.**
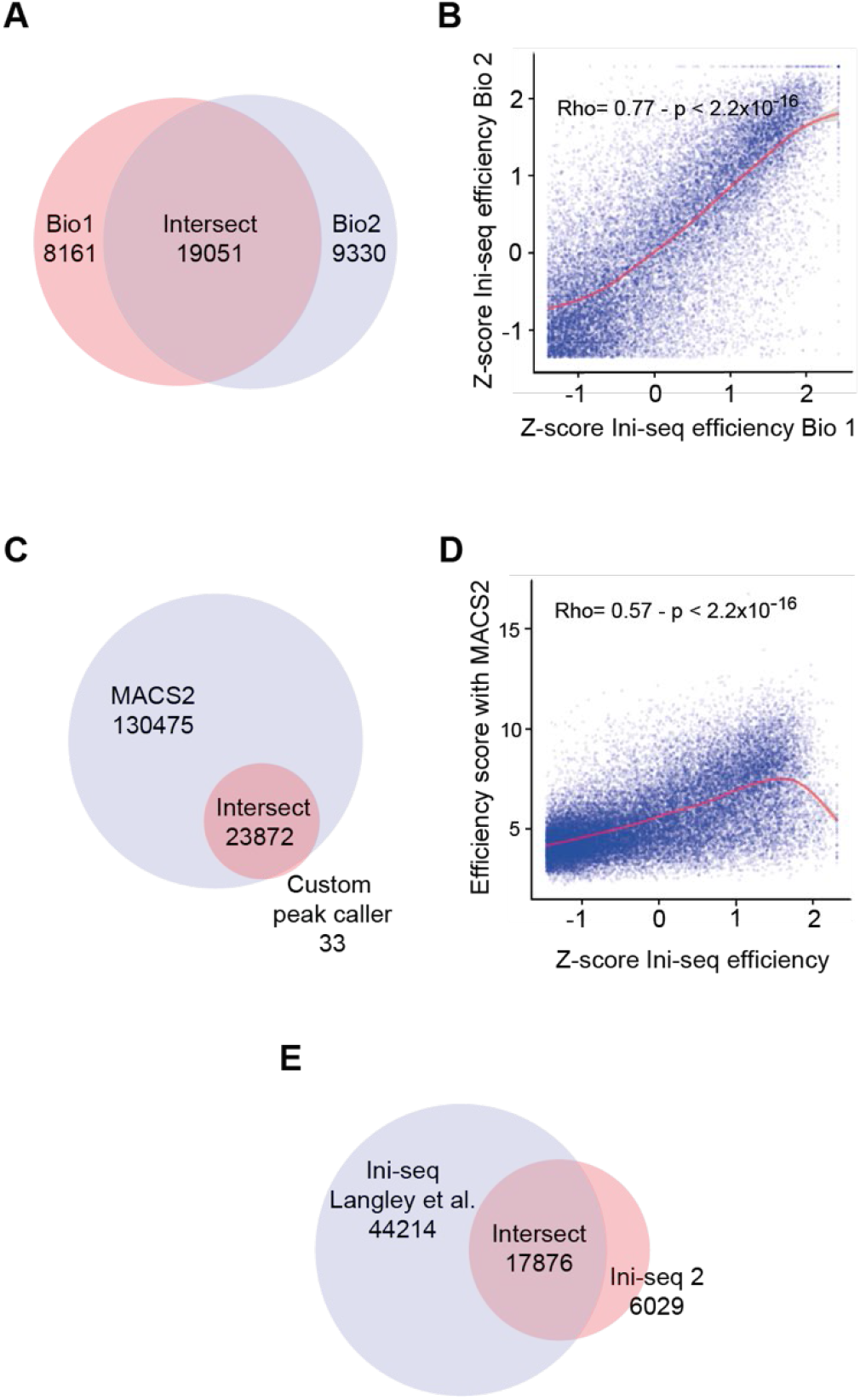
Biological replicates of ini-seq 2 and comparison between peak callers. A. Venn diagram of origins called in the two biological replicate ini-seq 2 experiments (terms Bio1 and Bio2). Permutation test p = 0.0001, Z-score 1256. B. Correlation of origin efficiency in the intersect of the biological replicates. Correlation: Pearson. C. Venn diagram of origins called by the custom peak caller described in this paper compared with MACS2 (1) using the parameters of Akerman et al. (2). Permutation test p = 0.0001, Z-score 538. D. Correlation of origin efficiency in the intersect of (C). Correlation: Pearson. E. Venn diagram of origins called in ini-seq 2 compared with the larger biological replicate of the original ini-seq experiments (3). Permutation test p = 0.0001, Z-score 286.

**Figure S2.**
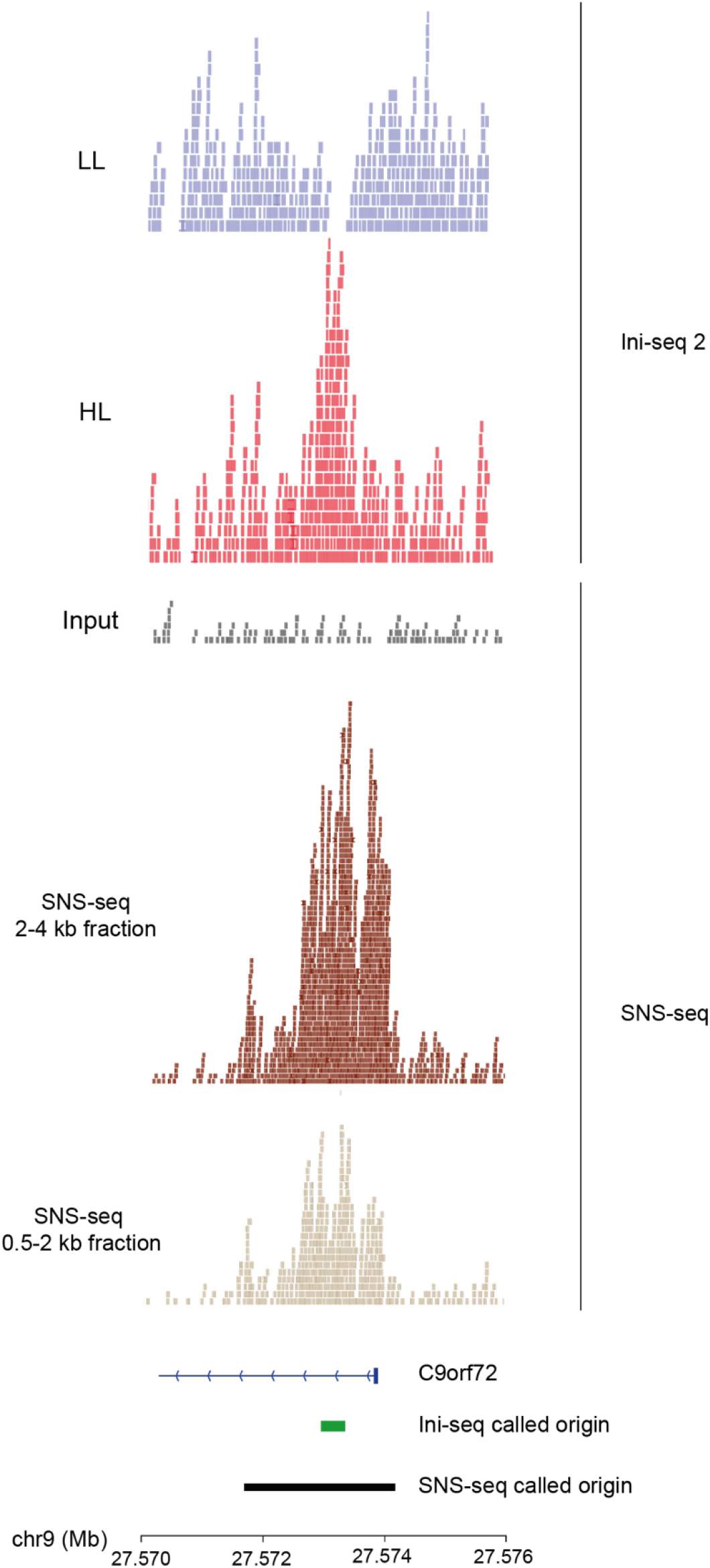
Example IGV tracks comparing ini-seq 2 with SNS-seq in EJ30 cells. A region at the start of the C9ORF72 locus on chromosome 9 is shown. From top to bottom: Ini-seq 2 light/light and heavy/light fractions with input (total genomic DNA) used for SNS-seq; SNS-seq reads from size fractions 2 – 4kb and 0.5 – 2 kb. The ini-seq 2 peak called with our custom peak caller is shown as a green horizontal bar; the SNS-seq peak called with MACS2 is shown in black.

**Figure S3.**
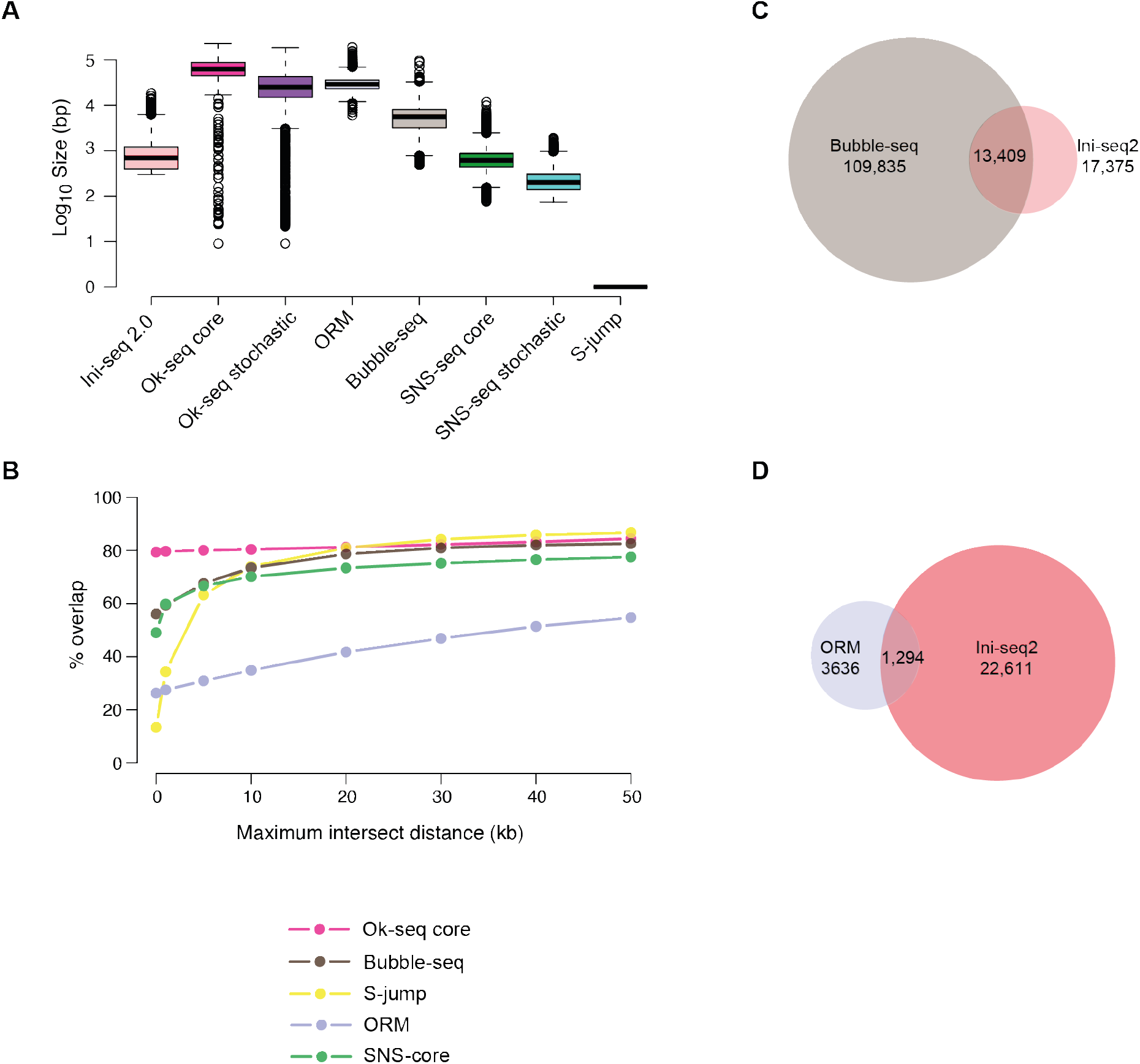
Further comparison of ini-seq 2 with other origin mapping techniques. A. Size distributions of origins called by the indicated different techniques. Whiskers represent interquartile range. B. Intersect between ini-seq 2 peaks and peaks called by other techniques as a function of maximum intersect distance allowed. The Y-axis gives the percentage overlap of the group with the smallest number of origins in each comparison. C. Venn diagram showing the overlap between ini-seq 2 origins and bubble-seq initiation sites (4). Maximum intersect distance allowed for overlap: 5kb. Permutation test p = 0.0001, Z-score 101. D. Venn diagram showing the overlap between ini-seq 2 origins and optical replication mapping (5). Maximum intersect distance allowed for overlap: 0kb. Permutation test p = 0.0001, Z-score 35.

**Figure S4.**
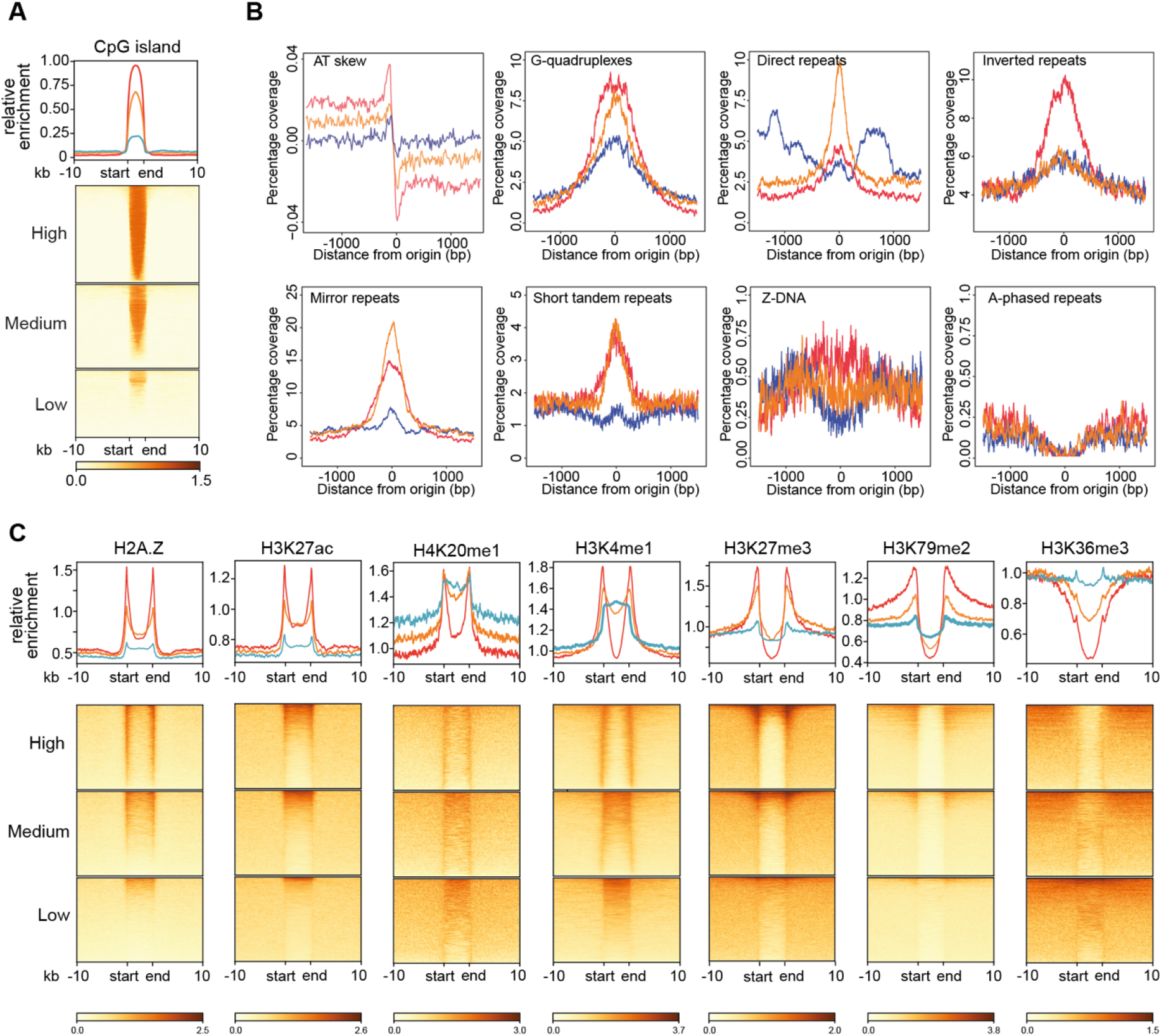
Genetic and epigenetic features of ini-seq 2 replication origins. A. Relative enrichment of CpG islands within and +/− 10kb around the ini-seq 2 origin classes of low (blue), medium (orange) and high (red) efficiency. The origin itself is represented as a metagene by ‘start’ and ‘end’. B. Distribution of AT skew, G quadruplex-forming sequences, direct repeats, inverted repeats, mirror repeats, short tandem repeats (2 – 6 bp repeat), Z-DNA and A-phased repeats (6) around origins of the low (blue), medium (orange) and high (red) efficiency classes. Areas of +/− 1200 bp of the ini-seq 2 origins are shown. C. Relative enrichment or depletion of H2A.Z, H3K27ac, H4K20me1, H3K4me1, H3K27me3, H3K79me2 and H3K36me3 within and +/− 10kb around the ini-seq 2 origin classes of low (blue), medium (orange) and high (red) efficiency. The origin itself is represented as a metagene by ‘start’ and ‘end’.

**Figure S5.**
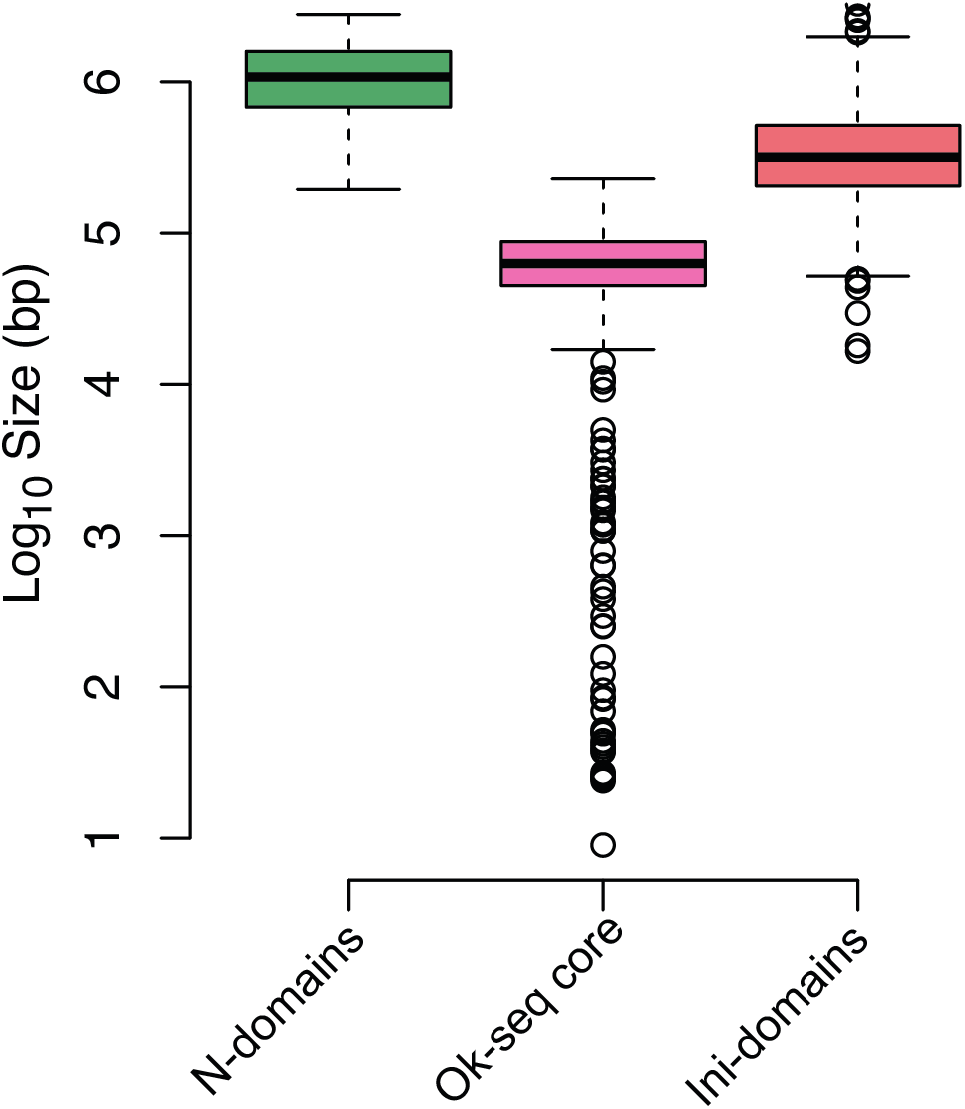
Size distributions of N-domains, Ok-seq core initations zones and ini-domains. Whiskers represent interquartile range.

## Notes

https://github.com/Sale-lab/Ini-seq-2

### Competing Interest Statement

The authors have declared no competing interest.

